# word2brain

**DOI:** 10.1101/299024

**Authors:** Abraham Nunes

## Abstract

Mapping brain functions to their underlying neural substrates is a central goal of cognitive neuroscience. Functional magnetic resonance imaging (fMRI) has proven indispensable in this endeavour. Recently, there has been growing interest in tackling this problem by mapping semantic concepts onto brain regions using repositories of images and text from the neuroimaging literature. However, no study has thus far approached this problem using (dense) vector representations of words. Using data from the Neurosynth database, we sought to develop a model that could (A) capture local correlations between words in text, as well as topics, (B) capture representation of distributed brain networks in relation to word embeddings, and (C) generate synthetic images given word inputs. We show that jointly embedding words and brain imaging data on a vector space can yield semantic representations that sensibly relate concepts across biological, psychological, and observational levels of analysis. Moreover, our proposed model makes no assumption about spatial orientation of fMRI voxels, which allows for embedding of distributed brain networks onto the semantic space. We demonstrate this capability by generating synthetic brain activation vectors from word inputs. Our model has the potential to advance neuroimaging meta-analysis as well as contextual word-embedding methods more broadly.

## 1 Introduction

The clinical neurosciences are relatively underdeveloped compared to specialties such as cardiology. The most important limiting factor is arguably that we have no working model of how psychological and behavioural phenomena are generated by underlying neural computations. Cardiologists can listen to the heart or look at imaging and identify the patient’s likely symptoms, and then tailor therapy or design research trials based on physiological reasoning. This is made possible by considering the circulatory system as an instance of a fluid-dynamical system driven by a pump. Since the heart’s function is mechanical, then observations of its mechanics can be directly input into a doctor’s internal model of the patient’s circulatory system, and consequences accurately predicted. But what of the brain? We know it is an information processing device, but a generalizable and consistent conceptual model that links neural function to behavioural output has remained elusive for all but the most simple motor and sensory functions.

Functional magnetic resonance imaging (fMRI) has provided insights into the putative roles many brain regions; while some appear to serve relatively specific roles [Levy and Glimcher, 2012], others are ostensibly involved in many behavioural phenomena and illnesses [Goodkind et al., 2015, Stalnaker et al., 2015]. This mixture of redundancy and specificity have challenged our ability to develop coherent mechanistic models of brain function. Notwithstanding these problems, neuroscience holds as a core assumption that some mapping from neural activation space to that of behavioural function exists [Churchland and Sejnowski, 1992].

The difficulty in finding this mapping may be due in part to the constraint that individual studies necessarily approach a narrow aspect of brain/psychological function using subdomain-specific tools and terminology. Each fMRI research study thus provides a small amount of data to a sparsely populated corpus of cognitive neuroimaging literature. For this reason, it has been of great interest to pool data for meta-analysis. One such effort has been that of the Neurosynth database [Yarkoni et al., 2011], which has automated the extraction of fMRI activation maps from published literature, and linked these to respective studies’ abstracts. More specifically, each of the >11,000 aggregated studies in the Neurosynth database has text from the study’s abstract, as well as *xyz* coordinates (in a standardized reference space) of fMRI activations. Such data are an important resource for the pursuit of a mapping between neural and psychological spaces.

Indeed, the Nerosynth database begs the question of whether semantic content of research abstracts can be mapped onto brain imaging data from the respective studies. Several authors have approached this question, and we review their work in Subsection 2.1. We will show that there remain two limitations in the employed methods. The first is that the language models employed have been count based, rather than based on vector representations. The second is that these models have not been able to model brain networks (which are spatially distributed), jointly with the semantic representations.

Thus, the present study seeks to learn a vector-representation of semantic concepts in the Neurosynth abstract database. In Subsection 2.2 we review a popular method for learning vector representations of words. In Subsection 2.3 we present a simple method for combining the vector-representations of words with neuroimaging data such that (A) a joint neural-linguistic vector representation is learned that (B) allows for representation of long-range brain-network activations in the semantic embedding space. We show that our method generates a sensible semantic embedding that sensibly relates concepts across biological, psychological, and observational levels of analysis.

## 2 Background

### 2.1 Previous Approaches to the Neurosynth Dataset

Neurosynth was introduced by Yarkoni et al. [2011]. It is an open database summarizing data from >11,000 fMRI studies. While there are many aspects to the Neurosynth repository, we focus on two elements. First, each study in the database is indexed by a PubMed identification number, which facilitates access to the study’s published abstract. Second, Neurosynth has parsed original articles for each study and extracted brain activation coordinates (corresponding to individual voxels in standardized reference spaces).

There have been several approaches to modeling the relationship between sudies’ brain images and the language in their scientific abstracts. The first notable approach was that of Poldrack et al. [2012], who applied Latent Dirichlet Allocation (LDA; Blei et al. [2003]) to the abstract data and then correlated the inferred topic sets with the neural activation coordinates of constituent studies. In LDA, one assumes each document is a distribution over topics, and that each document is a distribution over words. Poldrack et al. [2012] first applied LDA to the text topics the corpus. To determine the association between topics and voxels, the authors performed a chi-square test for each voxel in a given study against a binary indicator of whether that study loaded onto a given topic. This model measured association between topic loading and voxel activation, but in the process necessarily lost generative capacity.

An important limitation of this study was that they did not model activations jointly with the text. To address this, Rubin et al. [2016] later introduced Generalized-Correspondence Latent Dirichlet Allocation (GC-LDA), in which the topics in an LDA were defined as elliptical volumes in the brain (in the same *xyz* coordinates as the reported activations). The GC-LDA model thus defines brain volumes as probability distributions over words (i.e. those in abstracts of studies reporting neural activation coordinates), and documents as probability distributions over brain areas. Using deep restricted Boltzmann machines, Monti et al. [2016] also address this limitation by jointly modelling word counts with brain activation vectors.

Although the GC-LDA and restricted Boltzmann machine approaches tackled the joint modeling of neural activations and psychological phenomena (insofar as the latter can be captured by words of abstract text), there are several important limitations. Both models—by virtue of using total word count— could not capture local context of individual words in an abstract. This is important because there may be individual semantic meanings found in different local contexts within an abstract. In addition, LDA is limited in its ability to capture correlations between topics, owing to the nature of the Dirichlet prior. This would, for example, limit our ability to note the high correlation between topics such as eating-related behaviours and addiction [Gearhardt and Potenza, 2013, Volkow et al., 2013, 2017], unless they were included within a single topic. GC-LDA inherits LDA’s limitation with correlated topics. However, in this case it would preclude modeling of brain networks, since topics in GC-LDA are defined as contiguous (Gaussian) volumes. This is a significant limitation since it is likely that most psychological and behavioural functions of interest involve network-level computations [Petersen and Sporns, 2015, Sporns, 2013].

### 2.2 The GloVe Algorithm and Vector Representations of Words

Vector representations of words offer an important advantage over the whole-document bag of words models implemented in the prior approaches to Neurosynth. Specifically, vector representations model words in terms of how they are distributed in the context of other locally occurring tokens. This is particularly important for our domain of analysis. For instance, one expects to encounter the word “drug” often in neuroimaging literature, but its meaning will vary depending on whether it is in the context of the words “[drug] addiction” or “clinical [drug] trial.” The traditional topic models would suffer particularly in cases where a drug (medication) trial was being conducted for treatment of (illicit) drug addiction. Moreover, since neuroscience is a highly interdisciplinary field, one would expect to encounter multiple semantic concepts within a single abstract. While a detailed review of vector representations of words is beyond the scope of our paper, we focus on the *Global Vectors for Word Representation* (GloVe) model [Pennington et al., 2014], which we employ herein.

The GloVe model was introduced by Pennington et al. [2014] by the justification that word co-occurrences provide the most meaningful source of information for unsupervised learning. Rather than implementing learning at each step of movement of a sliding window over the corpus, as in the word2vec models [Mikolov et al., 2013a,b], the GloVe model tallies the co-occurrence of words up front (within windows of a fixed size). It is with these co-occurrence tallies that the GloVe model’s word vectors are subsequently fit. Due to space constraints, we present only the highlights of their model relevant for implementation (the reader is directed to the original paper for full details and an interesting analysis).

At the center of the GloVe model is a word co-occurrence matrix 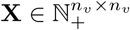 where *n*_*v*_ is the size of the vocabulary.

The entry *x*_*ij*_ of **X** denotes the number of times word *i* was encountered in the context of word *j*. We denote the set of all contexts in the corpus as *C*, the set of all contexts of word *j* as *C*(*j*), and a single *instance* of a context of word *j* as *c*(*j*). The co-occurrence matrix **X** is thus computed as

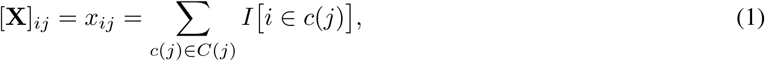

where ***I***(·) is an indicator function taking a value of 1 if *i* is in the specific context *c(j),* and 0 otherwise. Letting 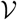 be the set defining our vocabulary, the total number of words co-occurring with *j* is thus

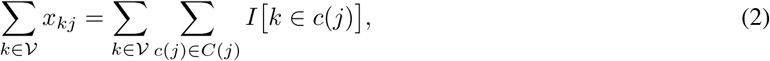

and the probability of encountering word *i* given *c*(*j*) is

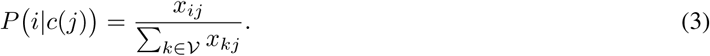

Since the matrix **X** is symmetrical, we can simplify our notation by stating that *P*(*i*|*c*(*j*)) = *P*(*i*|*j*). Now considering two different words *i* and *k,* each of which may occur in the context *c*(*j*), we can define a ratio of the posterior probabilities of *i* and *k*, given *j*:

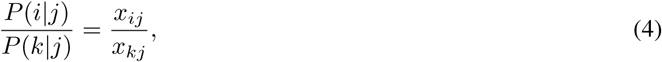

which should be large if *i* is semantically more similar to *j* than *k*, and small when the opposite is true. In cases where *i* and *k* are semantically similar, then this ratio should approach 1. Pennington et al. [2014] define the goal of learning a parameterized function *f* (·) mapping from a space 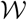 of latent parameters to the space of these word co-occurrences. For the triad *i, j, k* from Equation 4, Pennington et al. [2014] specified the following general model:

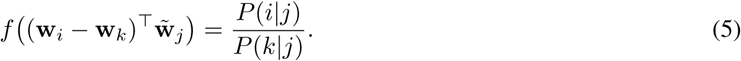

Assuming *f* (·) defines a structure-preserving mapping between the groups (ℝ, +) and (ℝ, ×) we can see that 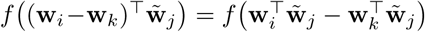 and *f* (·) can be further refined to

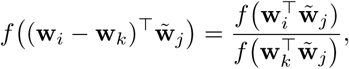

for which Pennington et al. [2014] identify a clear definition for *f* (·) as the exponential function. From this, we proceed to define the inner product term for *i* and *j*,

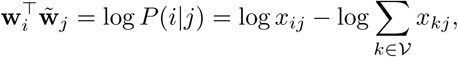

which can be made easier to work with by (A) accounting for the log-sum term in a bias term *b*_*i*_, and (B) adding a bias 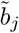 corresponding to the context parameters 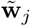. The resulting model is

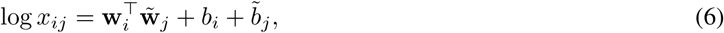

which is trained as a weighted least squares model under the following loss function:

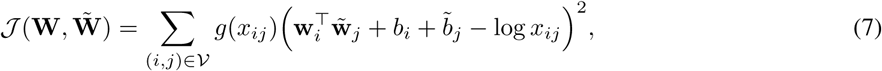

where **W** ∈ ℝ^*n*_*v*_ × *n*_*z*_^ and 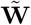 are the target word and context embedding matrices, respectively, and *n*_*z*_ is the dimensionality of the latent space. Pennington et al. [2014] implemented add-one smoothing to the logarithmic term and specified the weight function *g*(*·*) as follows:

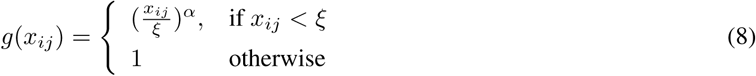

In Equation 8, the parameters *ξ* and *α* are specified *a priori*. Since some word pairings will co-occur far more often than others, one truncates their influence by clipping **X** at the value of *ξ*. The parameter *α* controls the curvature of the weighting function below *ξ*. Pennington et al. [2014] set these parameters to *ξ* = 100 and *α* = 3/4, which we also use in our analysis.

The strength of this model is that after training, the matrix **W** represents each word from the underlying corpus in a continuous *n*_*z*_ dimensional space. Perhaps more importantly is that a coherent measure of distance—the dot product—exists between words embedded in this vector space.

### 2.3 Embedding Words on Brain Activations

The GloVe method does not readily admit application to neuroimaging data. While one could forsee joint representation of neural activation coordinates in the space 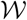, we propose a simpler approach.

The brain imaging data for the *d*^*th*^ study in the Neurosynth dataset is a 228,453 × 1 binary vector **m**^(*d*)^, where each unit is a voxel in the standardized Montreal Neurological Institute (MNI) space. We are interested in the degree to which words map onto individual voxels. We formulate this problem as follows.

We are given a word vector for document *d*, which is the sum of all embedded words in that abstract, 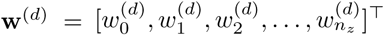 and a brain activation vector 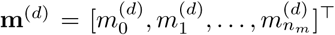. Since it is binary, we can view the outer product m^*(d)*^w^*(d)*⊤^ as effectively modeling a a “gating” effect of m^(*d*)^ on w^(*d*)^. That is, if element 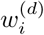 is nonzero, but is multiplied against an inactive voxel 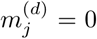 that component of the embedded word is nullified. Conversely, 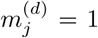 and 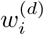 is large, and this pairing is encountered across many documents, that would suggest that the *j*^*th*^ brain voxel has a strong influence the *i*^*th*^ component of the word embedding space.

We compute an association matrix **C** ∈ ℝ^*n*_*m*_ × *n*_*z*_^ by summing these outer products over all studies

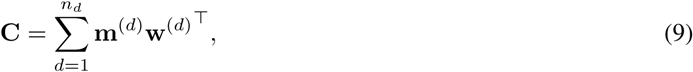

which we then submit to an (economical) singular value decomposition **C** = **USV**^⊤^. This SVD is economical in the sense that U ∈ ℝ^*n*_*m*_ × *n*_*z*_^ are the first *n*_*z*_ eigenvectors of **CC**^⊤^, **V** ∈ ℝ^*n*_*z*_ × *n*_*z*_^ are the eigenvectors of **C**^⊤^**C**,and **S** is a diagonal matrix with the square roots of the first *n*_*z*_ eigenvalues of both **C**^⊤^**C** and **CC**^⊤^.

While Equation 12 computes **C** using document vectors, this is essentially a weighting of **C** by the number of times a given word was used in the corpus. Formally, consider a corpus of size *n*_*d*_, where the *d*^*th*^ document consists of *n*_*t*_ ordered tokens (words) 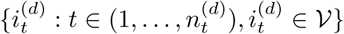 then each word’s corresponding embedded representation is 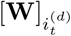 which we denote as 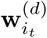 We define the document vector as

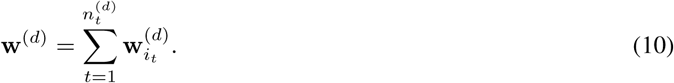

Substituting Equation 10 into 12, we therefore see that

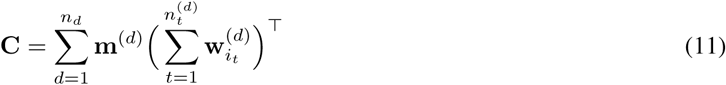

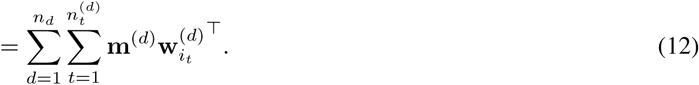

This approach is intuitively understood as involving two steps. First, the construction of **C** measures the movement of a vector in the word embedding space caused by activating a brain voxel (loosely speaking). In other words, one can think of brain activation patterns as warping the embedding space. One can view this warping as injection of real-world context into the semantic space. The second step of our approach is to calculate basis vectors for this joint neural-linguistic space, thereby “reshaping” it in a way that respects both the semantic relationshps between words, as well as those semantic relationships to the underlying substrate of their meaning: the brain’s activation patterns.

In addition to the advantage of being able to capture the local context of words through GloVe, our proposed model consists only of linear operations. Consequently, it becomes easy to map between word vectors, document vectors, and brain activation vectors. This facilitates interpretability and computational efficiency. Despite linearity in operations, our model also advances on previous approaches to the Neurosynth dataset by being the first to reasonably account for distributed network-level brain activations jointly with semantic word representations.

## 3 Methods

This section will guide the reader through the experimental workflow depicted in Figure 1. For parsimony, we describe details of the parameters for our implementations in Appendix A.

**Figure 1:**
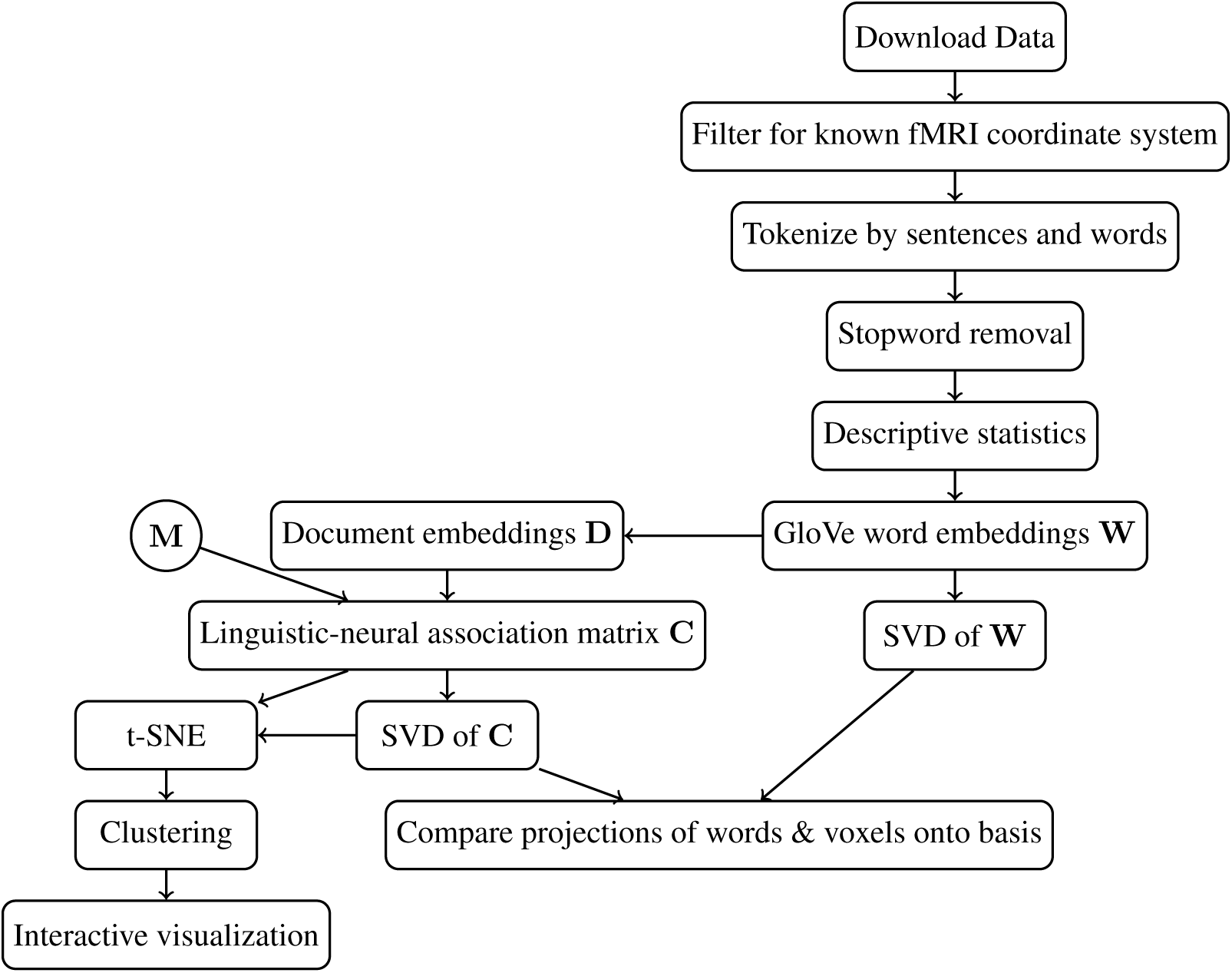
Methodological workflow. *Symbols:* association matrix **C**, which is the sum of outer products of each study’s neural activation vector and a document vector; neuroimaging activation matrix **M**, which carries a vector representation of the brain imaging data for each study; word embedding matrix **W**; matrix of document vectors (in rows) **D**. *Abbreviations:* functional magnetic resonance imaging (fMRI), singular value decomposition (SVD), t-distributed Stochastic Neighbour Embedding (t-SNE).

### 3.1 Dataset

Data were obtained using the neurosynth package^1^ for Python. Abstracts for 11,404 studies were downloaded. Preprocessing of abstract text was done using the nltk package in Python. Studies where neural activation coordinates were reported in an unknown space were removed (this included 1328 studies), leaving 10077 studies remaining. We first tokenized the abstracts into sentences, and then to words within sentences. Stop words and punctuation were removed, and text was converted to lower case. The original dataset included lists of authors; we removed names of the authors in cases where they appeared in an abstract of which they were an author. Other instances of names were left in place, since these may represent eponymous concepts such as “Wernicke” (an area of the brain, as well as a type of encephalopathy). Stopword and author name removal were done to reduce the dimensionality of word vectors, which are of the size of the vocabulary.

We computed descriptive statistics for these data, including total word, unique word, and sentence counts, as well as means and quantiles for these variables across studies. We summarized the total number of brain activations and the distribution thereof in a similar fashion.

### 3.2 Embeddings and Decompositions Thereof

For each of the 23,273 terms in the dataset, we created a co-occurrence matrix **X** ∈ ℝ^*n*_*v*_ × *n*_*v*_^, where entry *x*_*ij*_ represents the number of times term *i* was encountered in the context of term *j*. We defined contexts as windows of 5 words. This was implemented using the skip-gram function in the Natural Language Toolkit (nltk) package, which extracted all bi-grams within the defined 5-word window; the co-occurrences defined by these bigrams were then tallied into **X**. We then computed a vector embedding of words **W** ∈ ℝ^*n*_*v*_ × *n*_*z*_^—where *n*_*z*_ = 100 is the dimensionality of the embedding space—using the approach outlined in Section 2.2. Our implementation was built in TensorFlow v.1.5 [Abadi et al., 2015], wherein optimization of the loss function (Equation 7) was done with stochastic gradient descent using the ADAM optimizer [Kingma and Ba, 2015].

After obtaining word embeddings, we computed document vectors for each study (as shown in Equation 10) and concatenated these into a matrix **D** ∈ ℝ^*n*_*d*_ × *n*_*z*_. With this matrix and the *n*_*m*_ × *n*_*d*_^ matrix of brain activations M, we then computed the joint linguistic-neural association matrix **C** as shown in Equation 12.

The following singular value decomposition was performed on the word embedding matrix **W** (from the abstracts only):

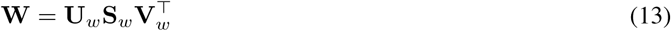

The first two components of **U**_*w*_ represent the projection of word vectors onto a 2-dimensional latent space defined by the SVD. For the joint linguistic-neural association matrix **C**, we performed the following SVD

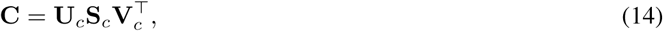

which defined a latent space onto which we projected **W** through the eigenvectors of the word embedding dimension, **V**_*c*_. The first two components of **U**_*c*_ define a mapping from the space of voxels to this 2-dimensional space.

### 3.3 Comparison of the Latent Spaces

We seek to compare how words segregate when projected onto a coordinate system defined only by those words, here **V**_*w*_, and when they are projected onto a coordinate system defined by both words and associated brain activity, here defined by **V**_*c*_. To do this, we evaluated how well words in **W** clustered when projected onto each of these respective spaces. This was done using hierarchical clustering with Ward linkage. We repeated this procedure while varying the number of clusters between 2 and 20. Since we had no labels for clusters *a priori,* the clustering performance was measured according to the silhouette score [Rousseeuw, 1987]. The result with the highest silhouette score was accepted as the final clustering, and a visualization of the stratified embedding presented.

#### 3.4 Generating Synthetic Brain Images from Text

We sought to evaluate the generative performance of our model by creating a function, `word2brain`, which takes a string of *n* words as an argument and generates a brain activation vector representing the hypothesized areas of activity given a description provided by those words. This was done by projecting the corresponding dense representation of each word supplied to the model onto the joint neural-linguistic embedding space, and subsequently projecting it back onto the *n*_*m*_ dimensional space of brain activation vectors. We visualize some examples using thresholded and unthresholded maps; the nilearn package [Abraham et al., 2014] was implemented for this purpose. The thresholded maps were plotted to determine whether key areas, such as the nucleus accumbens in addiction, could be identified. Unthresholded maps were plotted in order to examine the model’s ability to capture the general network associated with a given set of input words. The dimensionality of the joint embedding space was restricted to 10 (i.e. the first 10 principal components), although more or less could be used.

#### 3.5 Interactive Visualization

After performing the above analyses, we projected the document vectors **D** onto the space defined by the joint linguistic-neural representation, and subsequently ran t-Stochastic Neighbour Embedding [Van Der Maaten and Hinton, 2008]. The results were submitted to the hierarchical clustering procedure described above and visualized with the Bokeh package in Python [Bokeh Development Team, 2014]. An interactive version is available at http://www.abrahamnunes.com/plots/brainclusters/main.html, where the reader can explore how different studies are distributed along the joint linguistic-neural manifold.

## 4 Results

We began by evaluating the descriptive statistics of the Neurosynth dataset, which we present in Table 1. The vocabulary size in the present study was 23,273 and the total number of words in the corpus was 1,320,187. Each abstract had an average of 90 unique words (equal to median, 2.5th-97.5th quantiles=56.0-127.0; we use this quantile range henceforth). Each abstract had a mean of 8.8 sentences (median=9.0, 5.0-14.0), amounting to a total 88,669 sentences over the dataset. There were a total of 33,035,837 activated voxels reported in the dataset (i.e. number of non-zero elements of the matrix M). The average number of these entries for a given study was 3728.34 and ranged between 245.0 and 11440.5 (2.5% and 97.5% quartiles, respectively).

**Table 1:** Descriptive statistics for the Neurosynth Dataset. * Those studies reporting activations in an unidentified coordinate system were excluded (N=1328). *Abbreviations:* Montreal Neurological Institute (MNI)

|  | Over corpus | Study level statistics |  |  |  |
| --- | --- | --- | --- | --- | --- |
|  | Total | Mean | Percentile |  |  |
|  |  |  | 2.5% | 50% | 97.5% |
| **Studies** | 11,404 | - | - | - | - |
| **Studies Included\*** | 10,077 | - | - | - | - |
| **Activated Voxels** | 33,035,837 | 3728.34 | 245.0 | 2365.0 | 11440.5 |
| **Sentences** | 88,669 | 8.80 | 5.0 | 9.0 | 14.0 |
| **Words** | 1,320,187 | 131.0 | 74.0 | 131.0 | 192.0 |
| **Unique Words** | 23,273 | 90.0 | 56.0 | 90.0 | 127.0 |

### 4.1 Structure of the Embedding Space

The results of hierarchical clustering on words projected onto the principal components of **W** and **C** are presented in Figure 2. The best clustering performance was demonstrated by projection of words onto a space defined by the first two principal components of the joint neural-linguistic association matrix (silhouette score=0.46 for 2 cluster solution). We plot the projections in Figures 3, 4, and 5. These figures show that defining the semantic manifold with input from the brain activation vectors enhances the quality of semantic clusters on both quantitative (as per Figure 2) and qualitative grounds (as evinced through exploration of the embeddings). This is further supported by the data shown in Figure 6, which shows how the 100-dimensional axes in the original space of **W** project onto the first two principal axes of C, in comparison to their projection onto the principal components of **W** alone.

**Figure 2:**
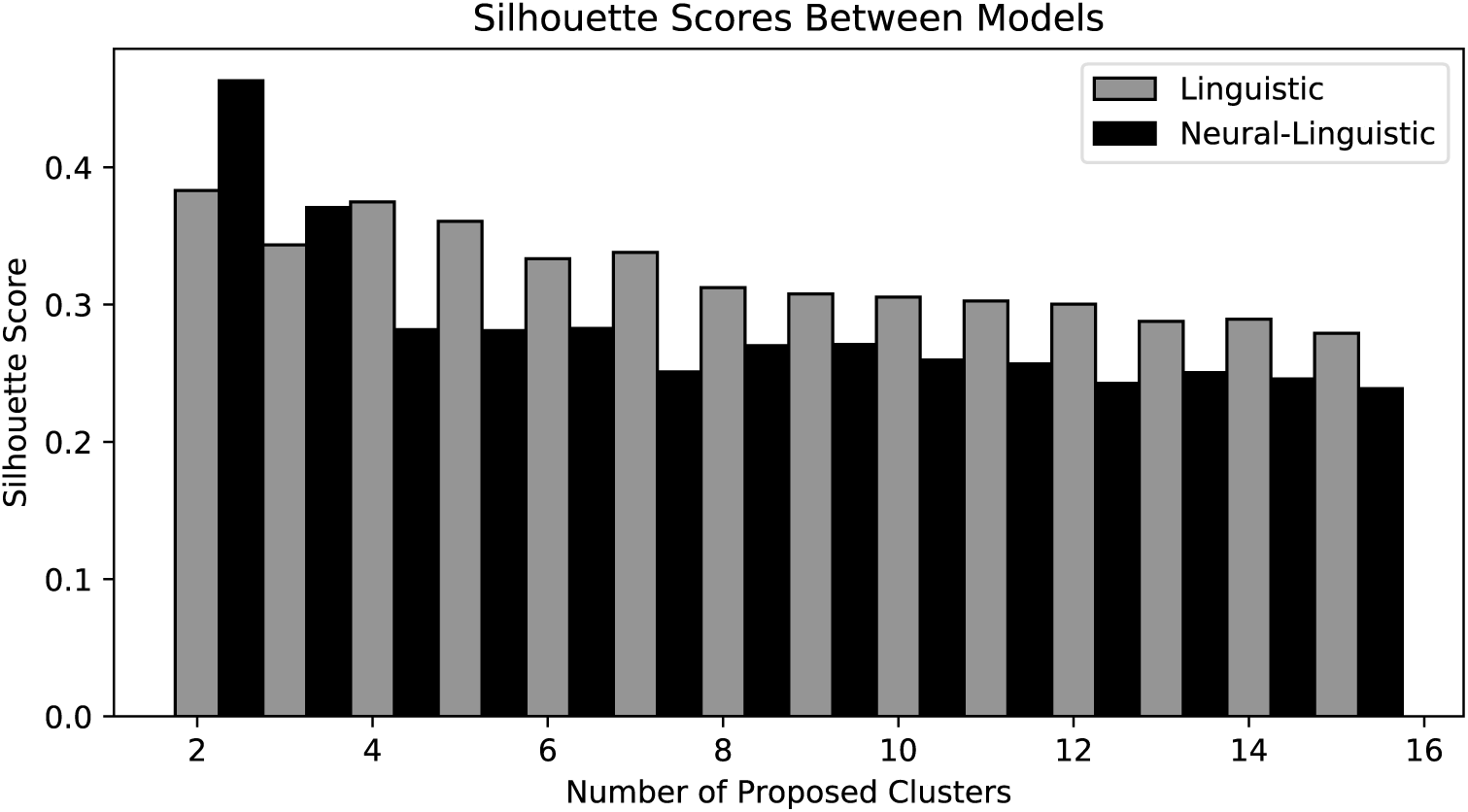
Silhouette scores computed for varying numbers of proposed clusters using (A) words projected onto a decomposition of the word embedding matrix **W** alone, shown with grey bars, and (B) a decomposition of the joint neural-linguistic association matrix, shown with black bars.

**Figure 3:**
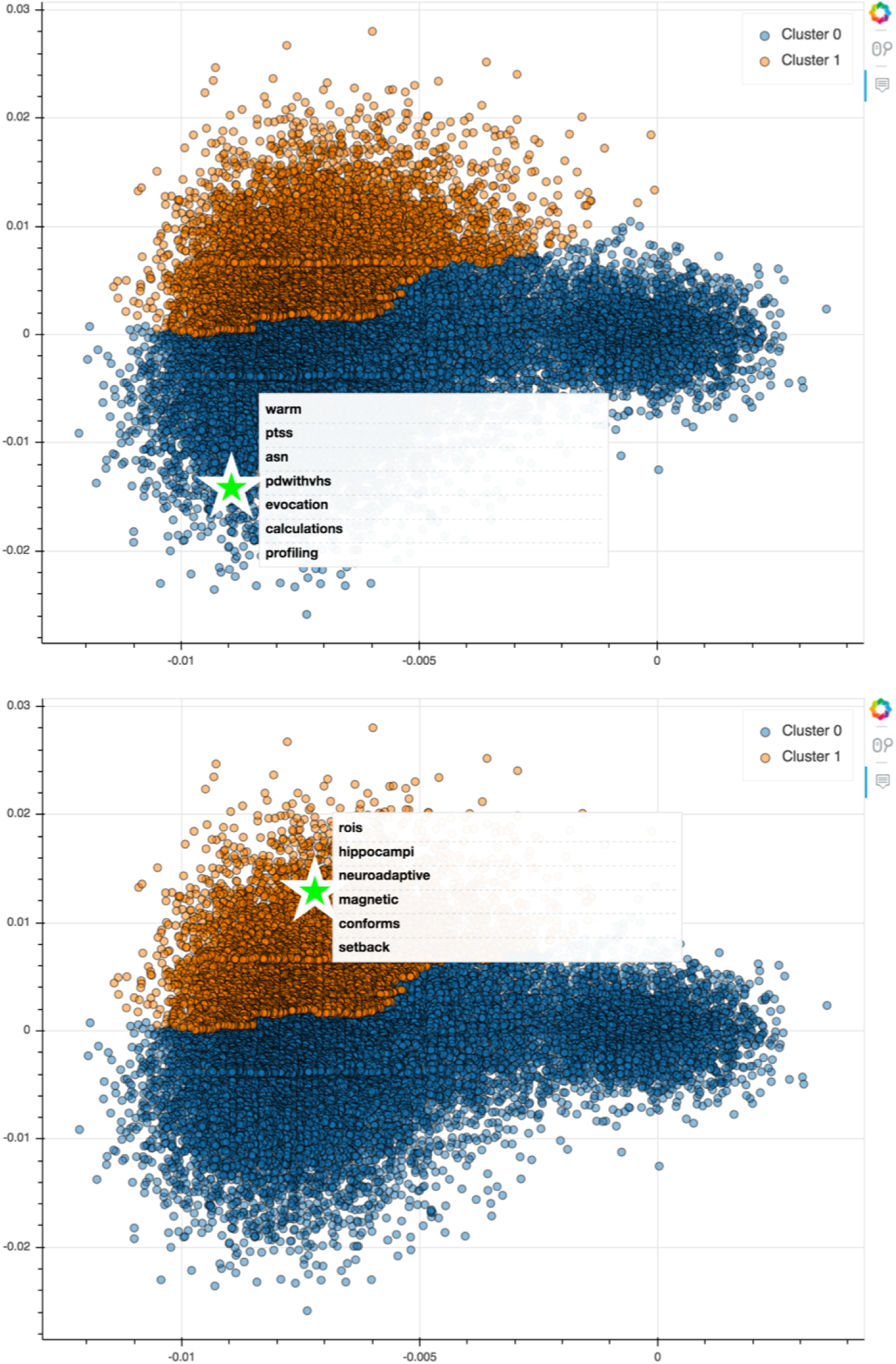
These are two plots of the projection of word vectors onto their first 2 principal components. Each point is a single word. The blue and orange colouring corresponds to the best-fit clusters on this space. We have highlighted the co-location of several words in the main clusters. Note that they lack semantic similarity, and also seem to have few brain related words. An interactive version of this visualization can be found at http://abrahamnunes.com/plots/brainclusters/main.html.

**Figure 4:**
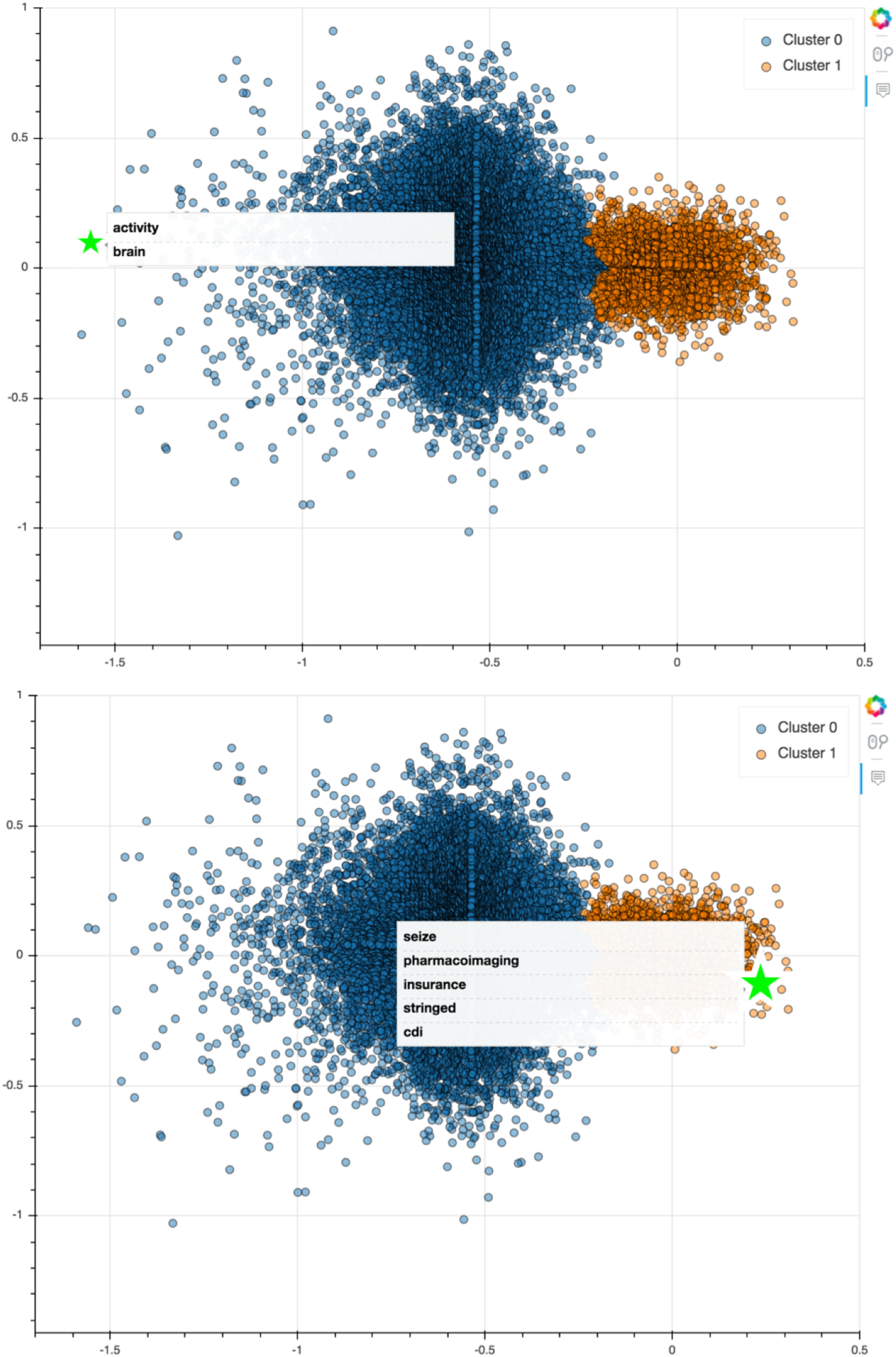
These are two plots of the projection of word vectors onto the first 2 principal components defined by the joint neural-linguistic association matrix **C**. Each point is a single word. The blue and orange colouring corresponds to the best-fit clusters on this space. We have highlighted the co-location of several words in the main clusters. Compare these results to those of Figure 3. Here, to the far left, we see that the two most extreme words in that direction are “brain” and “activity,” which are arguably the most obviously related words for fMRI research. Conversely, in the orange cluster, we see several words that are ostensibly unrelated to neuroimaging (although we show “pharmacoimaging” as an example here of one potential erroneous projection). http://abrahamnunes.com/plots/brainclusters/main.html.

**Figure 5:**
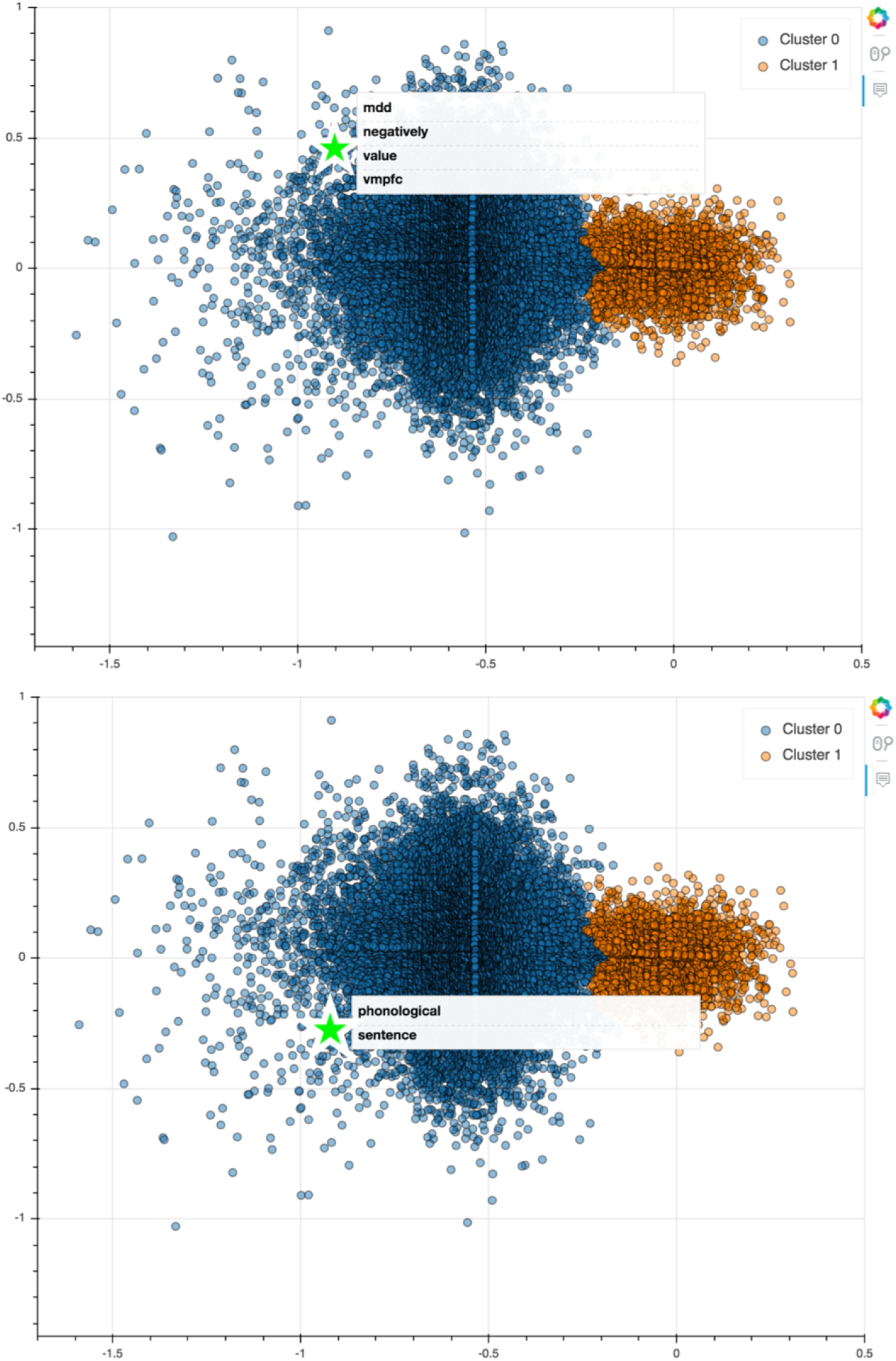
These are two plots of the projection of word vectors onto the first 2 principal components defined by the joint neural-linguistic association matrix **C**. Each point is a single word. The blue and orange colouring corresponds to the best-fit clusters on this space. We have highlighted the co-location of several words in the main clusters. Compare these results to those of Figure 3. Here, we see that our model has captured one of the most famous findings in functional neuroimaging, which is the relationship between the ventromedial prefrontal cortex (vmpfc) and value [Levy and Glimcher, 2012], along with major depressive disorder (mdd), and negativity (ostensibly negativity in value!). We also see that words related to language co-locate to the lower left. http://abrahamnunes.com/plots/brainclusters/main.html.

**Figure 6:**
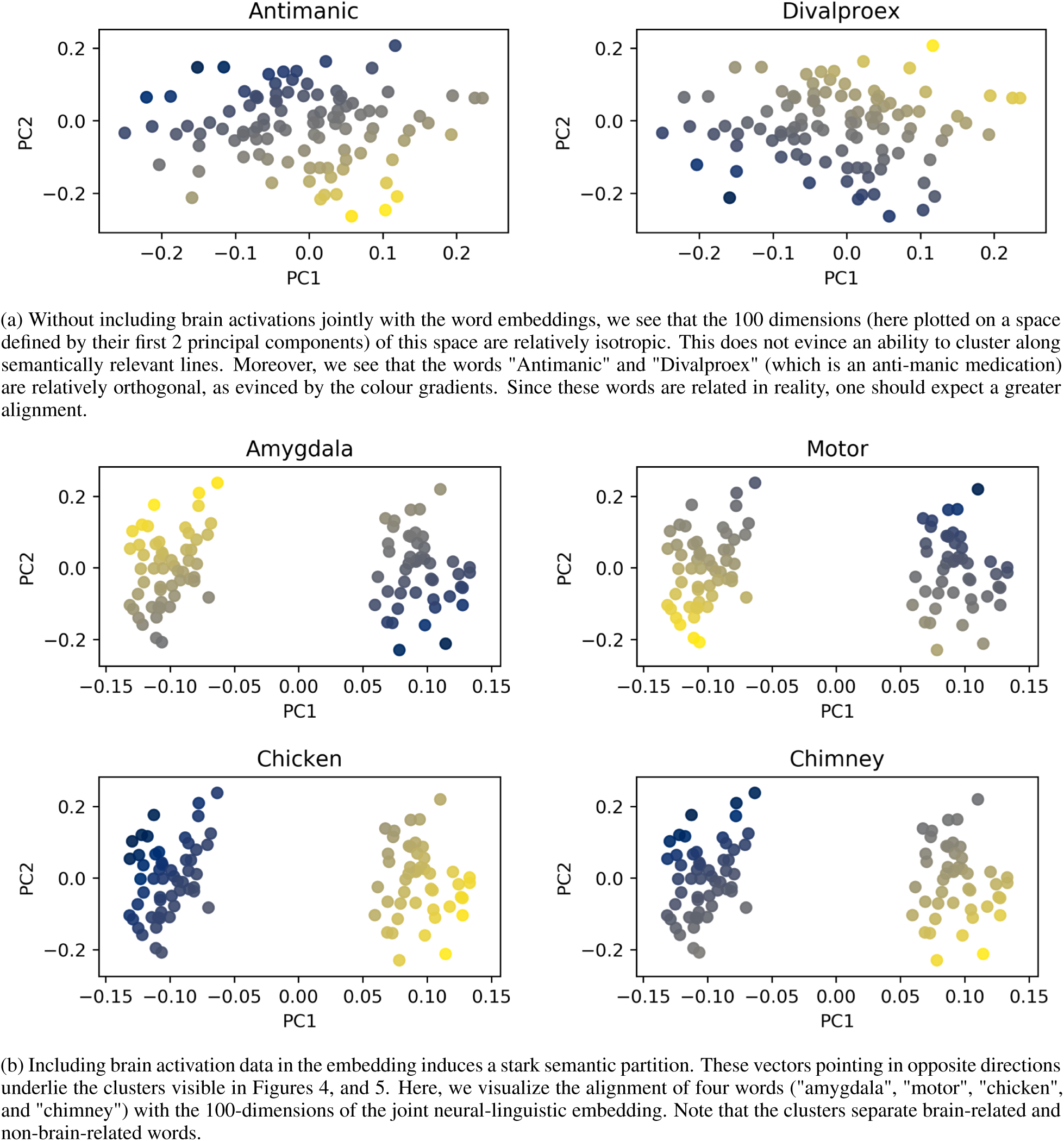
The principal components of the 100-dimensional embedding space. The uppermost row of plots represents the embedding space of words alone, whereas the bottom four plots show the space for a representation learned from decomposition of a joint neural-linguistic association matrix **C**. In each plot, we projected a word onto the defined space, and coloured each point according to how aligned that word is with that dimension of the embedding. Yellow indicates positive correlation, while dark blue denotes anticorrelation with a given direction.

The plot of Figure 6 indeed shows that the brain activation data reshape the word embedding space to segregate brain-related and non-brain related words. However, it is clear that beyond spreading along the first principal component, there is a further separation of semantic concepts along the second component. Moreover, this spread appears predom-inant in the brain-related words. To investigate this further, we projected (A) the brain activation matrix **M** and (B) the embedded words **W** onto the first two principal components of **C**. We plot each of the voxels as a point on the scatter plots of Figure 7. The projection of words onto this same space is represented by the dot colour, where red indicates that the word vector is oriented in the same direction as the voxels denoted by those points.

**Figure 7:**
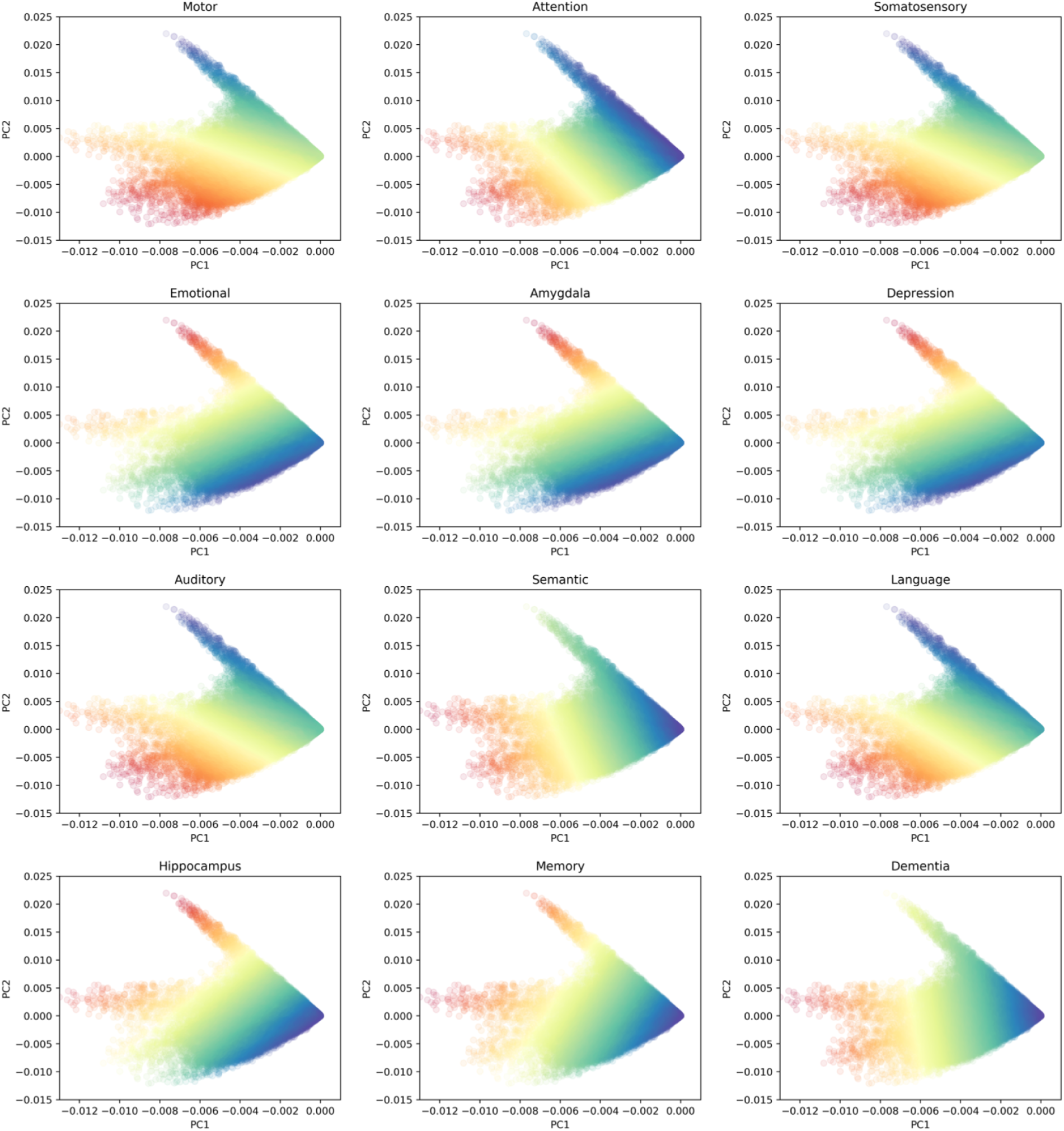
Projection of brain voxels onto the joint neural-linguistic space defined by the decomposition of **C**. Each plot shows the correlation of the projection of a word from our corpus onto this space. Note that the effect of the brain voxels is to further separate the semantic space ostensibly into emotional and unemotional domains.

**Figure 8:**
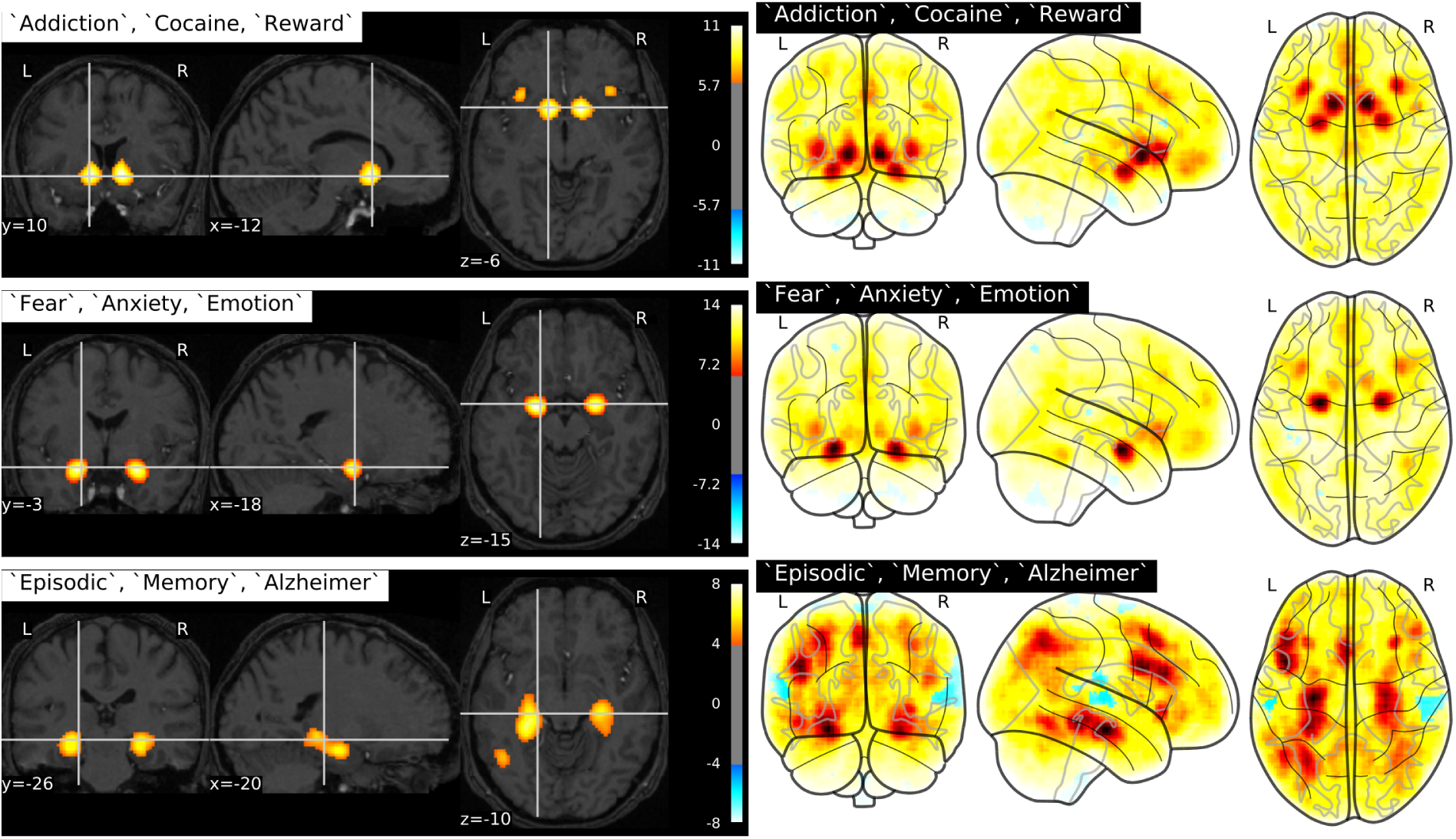
Activation maps synthesized from our model given the inputs of three words. The left column shows thresholded statistical maps centered at (from top to bottom) the nucleus accumbens, the amygdala, and the hippocampus. The right column shows unthresholded statistical maps, where deeper red indicates greater association, and blue rep-resents anticorrelation.

### 4.2 Generating Brain Images from Word Vectors

The data in Figure 7 suggest that some semantic mapping between word vectors and activity at brain voxels may exist. We therefore asked whether supplying a set of words to our model—that is, passing these words through the embedding space and onto the space of brain activation vectors—could generate statistical activation maps that correspond sensibly with known neurobiological relationships. Indeed, we found this to be the case. Some examples of these generated images are shown in Figure 9.

**Figure 9:**
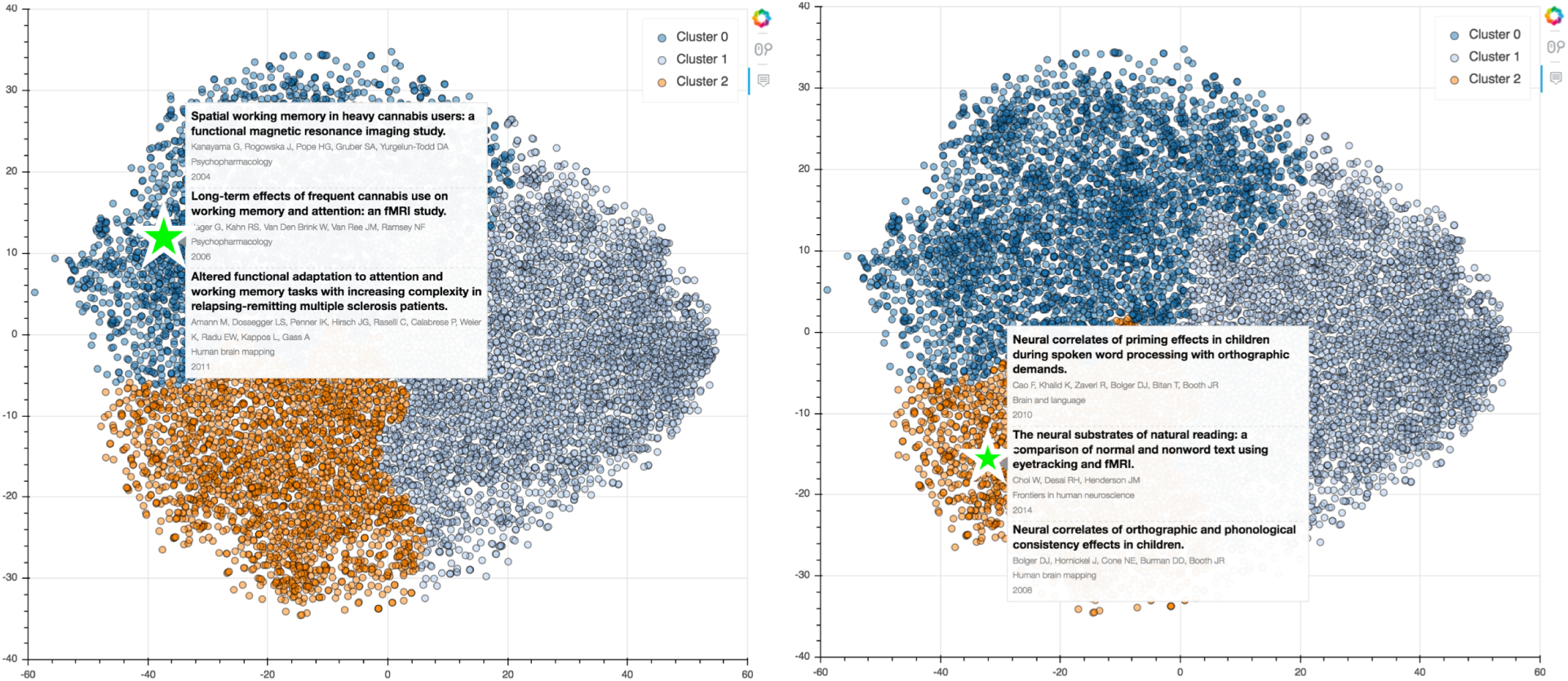
Examples of study coordinates on the joint linguistic-neural embedding space. Due to file size limitations, we show only two instances here. Further exploration of an interactive version of this plot can be done at http://www.abrahamnunes.com/plots/brainclusters/main.html.

### 4.3 Clustering of the Neuroimaging Literature

We projected the document vectors onto the joint neural-linguistic space, and further performed t-SNE in order to enhance visualization of those data. We show examples in Figure 9, and provide an interactive example at http://www.abrahamnunes.com/plots/brainclusters/main.html.

## 5 Discussion

We have shown that word embeddings combined with brain activation data can provide meaningful and interesting representations of semantic information in the cognitive neuroscience literature. Moreover, these representations are sufficiently powerful to generate hypothetical brain activation patterns given a set of input words. Our study makes important contributions both to research about the cognitive neuroscience domain, as well as to research on word embeddings.

First, we have shown that meaningful semantic relationships in neuroimaging research abstracts are suitable for vector space representations. This contributes to the body of knowledge for which Poldrack et al. [2012], Rubin et al. [2016], Alhazmi et al. [2017], Monti et al. [2016] and others have laid a foundation. Specifically, the existing approaches have been based on count based representations of words, which could not account for correlations between words in a local context, and in the case of LDA could not account for topic level correlations. In terms of the GC-LDA model of Rubin et al. [2016]—where topics were defined as brain volumes—this would preclude modeling network effects, since the topics were contiguous brain regions. While our GloVe-based approach bears some resemblance to latent semantic analysis [Pennington et al., 2014], which was applied to Neurosynth by Alhazmi et al. [2017], that study, too, considered a single document as a bag of words. Our method introduces one important difference (with respect to the language model), which is that a document is considered to be a sum of vectors in a vector space within which correlation between individual words can be captured.

There may be an interesting geometrical interpretation of why the vector space model works well in comparison to count-based methods. From the perspective of defining a manifold onto which neuroimaging activation vectors can be projected, the whole-document bag of words representation assumes that space to be separable along axes defined by each unique token. Geometrically, one can think about a set of *k* words defining *k* fixed search directions in some “true” semantic space Ω ⊂ ℝ^|Ω|^ with dimensionality |Ω|. We assume that brain activation vectors are—in the ground truth—projected by some mapping onto Ω, and the goal of analyses such as ours and those of Alhazmi et al. [2017], Poldrack et al. [2012], and Rubin et al. [2016] is, loosely speaking, to learn about these projections. We do this by evaluating the degree to which brain activation vectors and words co-align, with the assumption that this provides some information about Ω.

Unfortunately, it is unlikely *a priori* that a set of 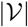 words, taken individually as fixed search directions, would form a positive basis for Ω if used in a count-based representation. If a count-based representation of 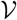 formed a positive basis for Ω, then any brain vector projected onto Ω would have a component fall in a half-space spanned by two word vectors, and thereby demonstrate some projection onto those words. For word-count representations, which are only meaningful if positive, we would would require at minimum |Ω| + 1 words to form a positive spanning set of search directions, and to be complete 2|Ω| words would be required (i.e. one word to search in a given direction, and another to search in the opposite direction). Conversely, with vector representations of words, anticorrelation is meaningful, which enables a single word to capture relevant semantic projections in Ω, bidirectionally. Finally, since vector representations of words can be correlated with each other and be of different lengths, there is a greater potential for defining complex search trajectories in Ω, and thereby improving the chances of capturing semantic information about brain activation patterns using linguistic data.

We have also shown that including real world context—that is, information about the actual subject matter of discourse—can improve the latent embedding space’s ability to capture relevant semantic structure. This was shown by the superior clustering performance with the joint neural-linguistic representation, compared to that observed with word embeddings considered alone. To our knowledge, this effect has not yet been shown in the literature. Given that the word embeddings were initially learned from the text data alone, our results do not suggest that inclusion of brain imaging data in the embedding process shows its effects on parameter optimization. Indeed, the same vector representations of words were used for both singular value decompositions in our analysis. What this suggests is that inclusion of the brain imaging data altered the shape of the manifold onto which the word vectors were projected. The resulting distributions of word vectors on this manifold were clearly superior in terms of both quantitative and qualitative measures of the sensibility of semantic clusters. More generally, our results may suggest that vector space models of semantics could benefit from including auxiliary data concerning the subject matter of the text from which those embeddings were learned. Future work should investigate whether this applies to the inclusion of multiple forms of auxiliary data; for analyses of Neurosynth, this may include merger of neurogenetic data from other repositories.

Our study is not free of limitations. First, we did not conduct a systematic analysis of hyperparameter settings for our models. This is largely due to the lack of clear guidelines or performance metrics that define a good embedding fit. Consequently, the hyperparameter tuning for these models is somewhat idiosyncratic. We opted to maintain consistency with previous literature, and indeed those choices performed satisfactorily. Future work should systematically explore the effects of hyperparameter tuning on GloVe embeddings. Another limitation is that we did not compare the joint linguistic-neural embedding against a latent representation of neural activations alone. This was omitted on account of the size of the document by voxel brain activation matrix, which precluded feasible implementation of singular value decomposition. These results would not have undermined the present study’s data, but could contribute to further understanding of how contextual information provided by the brain imaging data shape the embedding space; in future work we will consider application of matrix factorization approaches that facilitate subsampling in order to perform this comparison.

Notwithstanding these limitations, our study’s first strength is in the simplicity of our model. Being entirely linear, it enables mapping from word vectors to brain images, and has the potential for enabling automated captioning of brain images (an important target for future work). This carries significance for the future of neuroimaging meta-analysis. Current meta-analytic approaches rely on measuring some intersection of activations between studies that include some given linguistic term; our approach enables sampling of new hypothetical images from an underlying semantic space that incorporates information from the corpus of neuroimaging literature.

Our study’s second strength is the detailed analysis regarding the underlying model mechanics. Specifically, we were able to provide empirical rationales for both (A) the effectiveness, and (B) mechanism of action, of joint embeddings compared to word-embeddings alone. Future work should focus on development of measures for uncertainty of vector-space representations (i.e. probabilistic word embedding models), as well as methods to improve the quality of image sampling from the embedding space.

### 5.1 Conclusions

In conclusion, we have shown that a vector space representation of linguistic data from scientific abstracts of functional brain imaging research—when combined into a joint embedding with brain imaging data themselves—can offer a rich and sensible distribution of semantic concepts that span biological, psychological, and behavioural levels of analysis. Moreover, we have shown that these representations can (A) capture correlations between words and (B) adequately represent documents and higher order topics through simple arithmetic operations. Our analysis suggests that the joint neural-linguistic embedding approach functions because the real-world contextual data reshapes the manifold onto which linguistic vector representations are projected. This approach not only shows interesting clustering of words, brain voxels, and documents, but can also generate synthetic brain images that would be expected given a set of words. Future work should consider methods to optimize the hyperparameters of this approach, generate probabilistic embeddings, and incorporate multiple forms of biological data in the embedding process.

## Acknowledgments

Dr. Sageev Oore provided helpful questions and feedback during the brainstorming stage, regarding potential exten-sions to GC-LDA.

## A Experimental Parameters

**Table 2:**
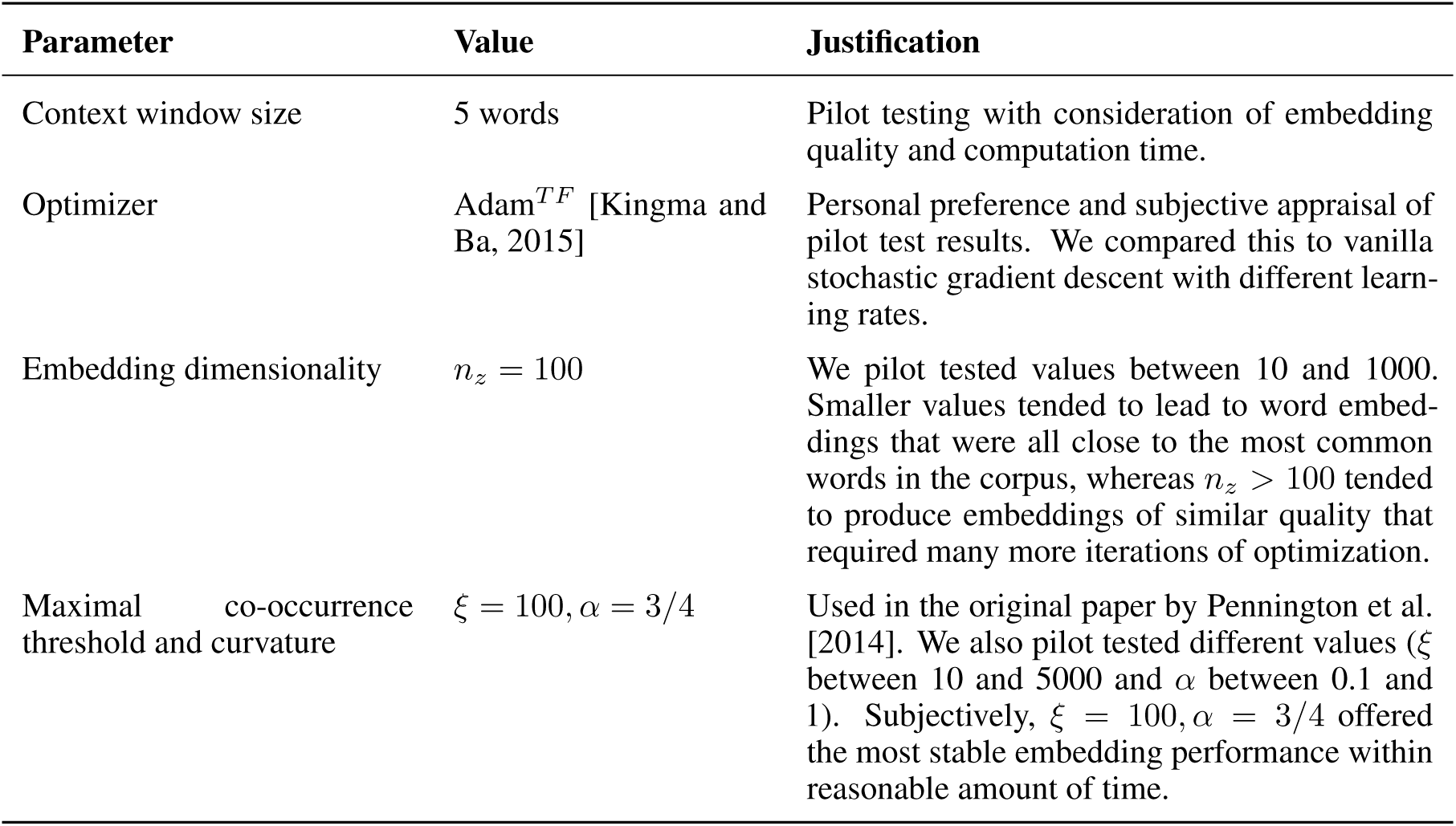
Experimental parameters and rationales. Items marked with ^*TF*^ were implemented in TensorFlow v1.4 [Abadi et al., 2015]. For pre-packaged implementations listed here, default parameters were used unless otherwise stated.

## Footnotes

https://github.com/neurosynth/neurosynth

